# Lake overturn as a key driver for methane oxidation

**DOI:** 10.1101/689182

**Authors:** M. Zimmermann, M. J. Mayr, D. Bouffard, W. Eugster, T. Steinsberger, B. Wehrli, A. Brand, H. Bürgmann

## Abstract

Many seasonally stratified lakes accumulate substantial amounts of the greenhouse gas methane in the anoxic zone. Methane oxidizing bacteria in the water column act as a converter, oxidizing methane into carbon dioxide and biomass before it reaches the atmosphere. Current observations and estimates of this methane oxidation efficiency are diverging, especially for the lake overturn period. Here we combine a model of turbulent mixing, gas exchange and microbial growth with a comprehensive data set for autumn mixing to quantify the relevant physical and microbial processes. We show that the microbial methane converter is effectively transforming the increased methane flux during the overturn period. Only rare events of pronounced surface cooling in combination with persistently strong wind can trigger substantial outgassing. In the context of climate change, these results suggest that changes in the frequency of storms may be even more important for methane emissions from temperate lakes than gradual warming.

## Main

Lakes and reservoirs are increasingly recognized as significant players in the global carbon cycle because they store, convert and release substantial amounts of greenhouse gases^1^. According to recent estimates, lacustrine methane emissions are responsible for about 75% of the climate impact of lacustrine systems^2^ and might even offset the continental carbon sink^3^. In lakes, most methane is produced in the anaerobic sediment as one of the final products of anaerobic carbon mineralization^4^. Ultimately, this methane is either released to the atmosphere or is oxidized by aerobic methane oxidizing bacteria (MOB), which have the unique ability to use methane as their sole carbon and energy source^5^. Here we provide evidence that this microbial methane converter is efficient and robust even at high rates of vertical mixing and prevents a major fraction of methane from reaching the atmosphere.

This microbial methane sink has important implications for lakes with seasonal thermal stratification that accumulate large amounts of methane in the anoxic hypolimnion, hereafter referred to as ‘stored methane’. During stratification, MOB at the oxic-anoxic interface form an effective barrier against diffusive methane outgassing, which is only bypassed by ebullition, i.e. methane bubbles released from the sediment^6^. In contrast, strong vertical mixing during the autumn overturn may lead to a rapid transport of stored methane to the surface. So far, estimates of global methane emissions from lakes have therefore assumed that most of this methane is ultimately released to the atmosphere_7_. However, the supply of methane to the oxygen-rich surface water creates favourable conditions for the growth of aerobic MOB. If the community of MOB grows fast enough, it could significantly reduce methane emissions even under conditions of rapid lake mixing.

Published field measurements of the percentage of stored methane that is oxidized range from 54 to 94%^8–11^. These previous studies focused either on time-resolved flux measurements^7,10^ or on microbial rate observations^8,9^. Here we closely integrate the two approaches in order to improve our understanding of the underlying processes during lake mixing. Based on a detailed field campaign we further developed and validated a dynamic model of turbulent mixing, gas exchange and Monod-type growth of MOB. We used this model to derive bulk kinetic properties of the MOB community and to quantify the efficiency and robustness of the microbial methane converter during the autumn overturn in a small temperate lake.

## Fate of methane during the autumn overturn

We determined the fate of methane in a small temperate lake during the autumn overturn (Figure 1). From October to December 2016 only 9% of the stored methane that was transported to the surface was emitted to the atmosphere. Methane emissions measured by eddy covariance flux measurement were on average 0.26 μg CH_4_ m^-2^ s^-1^ and summed to a total of 0.36 Mg C that were emitted from the whole lake (Figure 1a). During this period, cooling of the surface water as well as wind (Figure 1a) increased the mixed-layer depth from 7.6 to 13 m (Figure 1b). This physical process gradually depleted the pool of stored methane in the hypolimnion and transferred a total of 4.2 Mg C of methane to the mixed layer. After subtraction of the emissions, the remaining 3.8 Mg C either were oxidized by MOB or remained in the mixed layer. However, we only measured mixed layer methane concentrations of 0.5 ± 0.4 μM (SD) or a maximum total methane inventory of 0.039 ± 0.026 Mg C (SD), suggesting that MOB oxidized about 91% of the stored methane. In parallel, the cumulative biomass of MOB in the oxic mixed layer increased by a factor of 14 from 0.047 Mg C to 0.67 Mg C (Figure 1c). Despite this rapid growth, the 0.62 Mg C of newly formed biomass represented a small fraction of the 3.8 Mg C that were supplied to the mixed layer suggesting a carbon conversion efficiency of MOB of only 0.16. With the increase of MOB biomass we also observed a change in the community composition which is documented in detail in a parallel study^12^.

**Figure 1:**
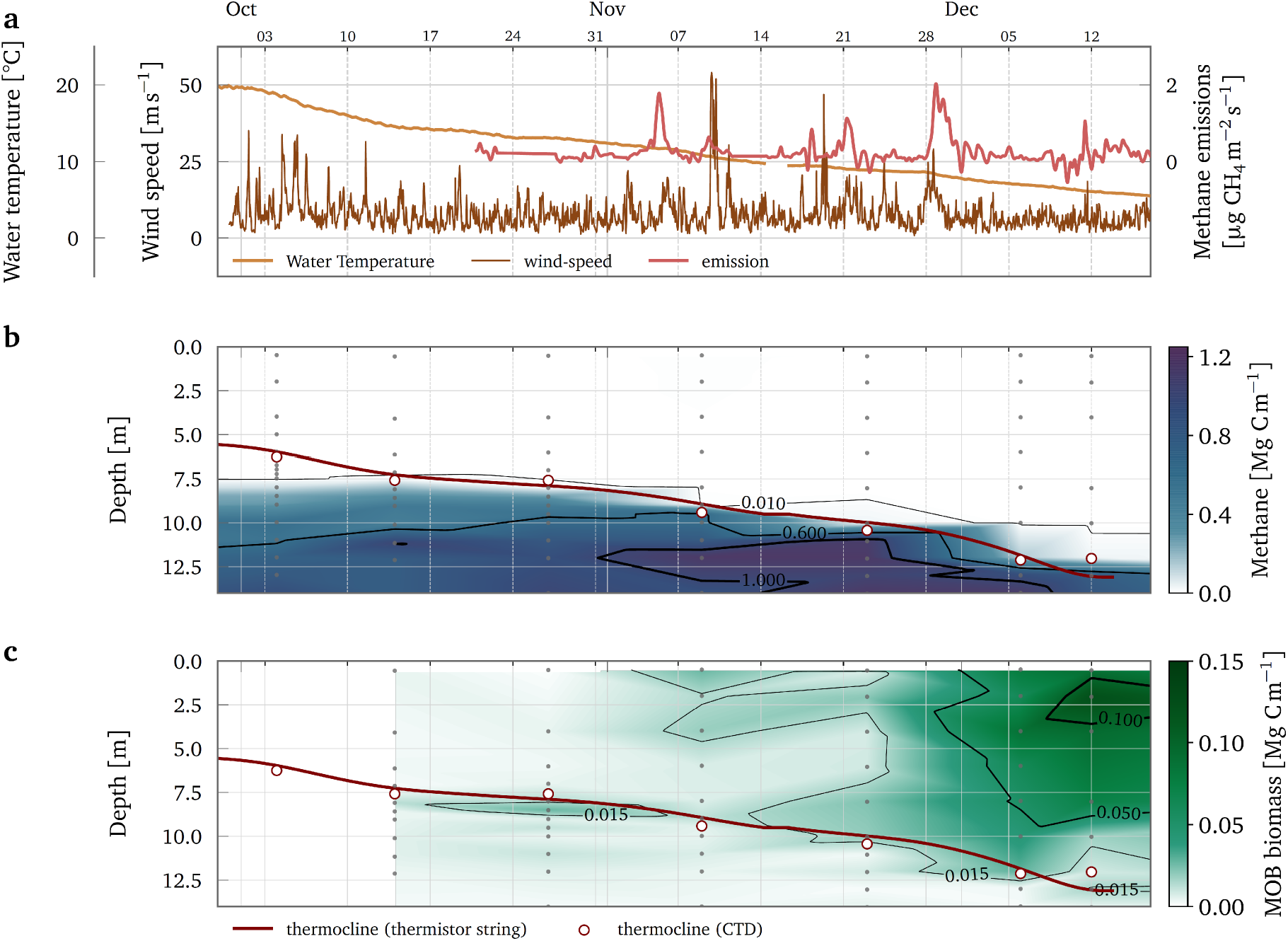
Field observations during the overturn period 2016 in Lake Rotsee. The horizontal time axis for all panels is plotted on top. Major ticks indicate the start of the month and minor ticks refer to the dates. (**a**) The daily average difference between air temperature and surface water temperature is plotted as a proxy for the progressing cooling during the overturn period. The daily maximum and minimum difference are indicated by the shaded area around the daily average. Wind-speeds are plotted as hourly averages and were corrected for Lake Rotsee (Supplementary Figure 1). Time series of the daily average air temperatures and wind-speeds were obtained from the closest automated weather station of MeteoSwiss (www.meteoswiss.ch). Methane emissions are plotted as Gaussian filtered flux of the measured 30 min average fluxes. (**b**) Methane stored in the water column multiplied by the cross-section area is plotted as a heatmap interpolated from individual measurements that are indicated by gray dots. Methane concentrations in the mixed layer were 0.5 ± 0.4 μM (SD) but reached levels of up to 3.7 mM at the sediment-water interface (Supplementary Figure 2), which corresponds to a partial pressure of about 65% at this depth. The depth of the thermocline was interpolated from the continuous thermistor string data and was verified by individual CTD profiles. (**d**) Biomass of MOB multiplied by the cross-section area is plotted as a heatmap interpolated from individual measurements that are indicated by gray dots.

## Modelling the key processes

Based on these field observations we validated a minimal model of the key processes (Figure 2) that determine the fate of stored methane and control the efficiency and robustness of the microbial methane converter during the autumn overturn. Despite its simplicity, the turbulent mixing model was able to reproduce the evolution of the measured mixed-layer depth (Figure 2a) with a root mean square error (RMSE) of only 0.27 m. The resulting flux of stored methane to the mixed layer was therefore in good agreement with the measurements (Figure 2b). The Monod-type growth model reproduced the evolution of MOB biomass to a maximum of 0.68 Mg C with a RMSE of 0.061 Mg C (Figure 2c). The box model approach neglected a potential gradual crossover of methane and oxygen within the thermocline. However, the biomass profiles did not give any indication that such a transition zone was of particular importance for methane oxidation during the overturn period (Figure 1c).

The ratio of modelled methane emission to the total transport of stored methane to the mixed layer suggests that about 98% of this methane was oxidized by MOB. The measured cumulative methane flux the atmosphere was about three times larger than the modelled flux (Figure 2d). A likely explanation for the discrepancy is the contribution of methane ebullition to the emissions during the overturn period, which was not included in the model. That bubble emission is a major pathway during the autumn overturn in Lake Rotsee has been shown in a study by Schubert et al.^8^ who estimated cumulative contributions of ebullition to total atmospheric emissions between 88 and 97%.

**Figure 2:**
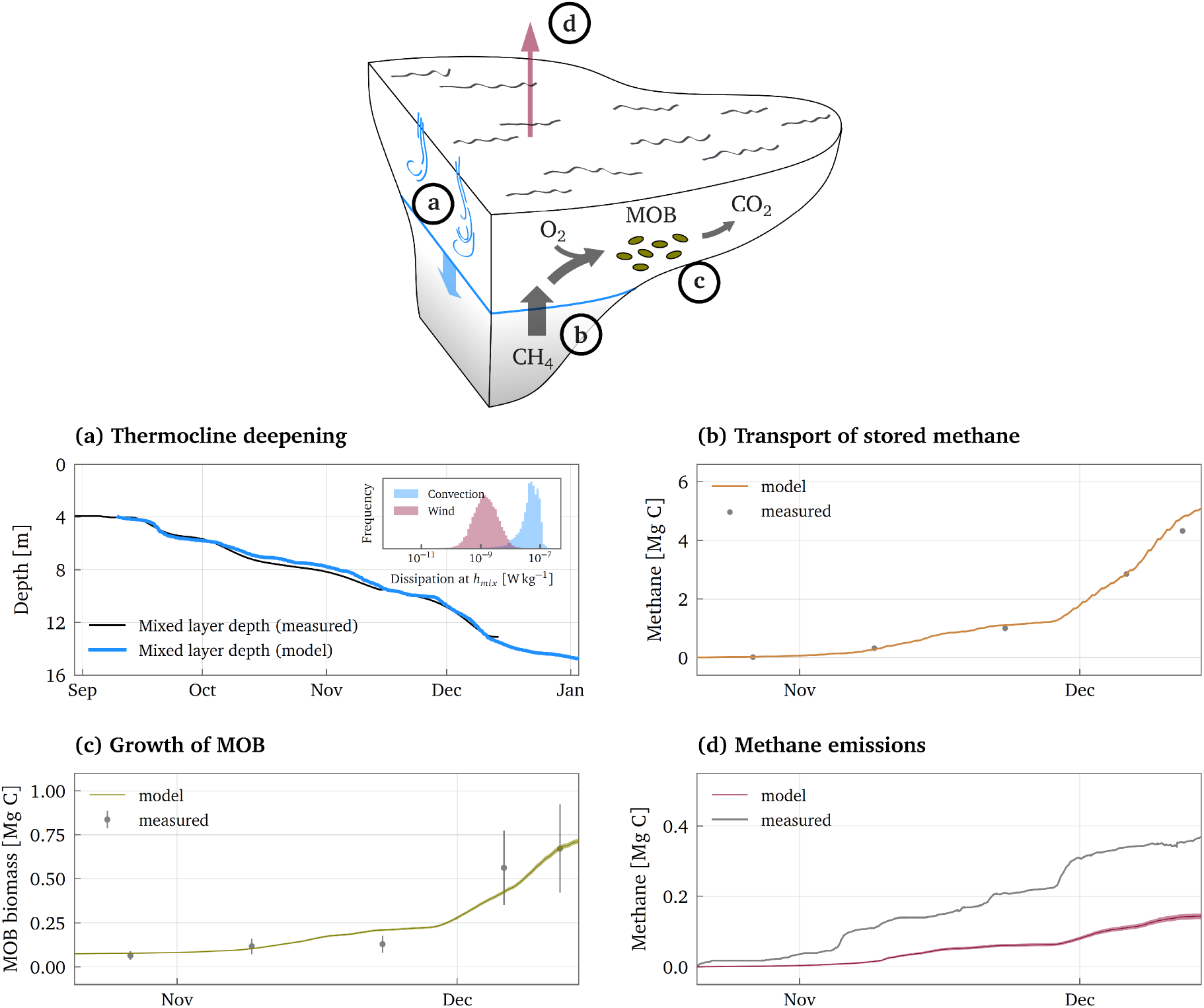
Model combining the key processes that determine the fate of stored methane during the autumn overturn. **(a)** The deepening of the thermocline was modeled using a turbulent mixing model that considers wind-driven and thermal convection. Modelled and measured depth of the thermocline fit very well. The inset shows the histogram of the dissipation rate of convective and wind-driven turbulent kinetic energy at the mixed-layer depth over the simulation period. **(b)** Modelled and measured cumulative amount of stored methane transported to the mixed layer. **(c)** Modelled and measured MOB biomass. The error bars for the measured biomasses represent 95% confidence intervals for the biomass, based on the uncertainty in the cellular carbon content. **(d)** Modelled and measured emissions of methane to the atmosphere. Emissions measured by eddy covariance also include ebullition, whereas the model does not include this emission pathway.

Even though the model does not take into account the observed diversity of the MOB community^12^, the fitted bulk kinetic parameter are well in line with expected properties for the methane limiting conditions in the mixed layer. The half saturation constant for methane oxidation (*K_M,CH_4__*) of 1.7 μM is in the lower range of published values (1 and 10 μM)^5,13^. It is known that the affinity for methane can even be in the nM range for growth under atmospheric methane concentrations^14,15^. The fitted half saturation in the lower μM range therefore reflects an adaptation to the low methane concentrations in the mixed layer. The best fit estimated for the carbon conversion efficiency of the Monod-type growth model (y) is 0.13 which is in good agreement with the 0.16 estimated from the measurements. Published values for the carbon conversion efficiency (CCE) of MOB vary widely from 0.19 to 0.7 and likely also depend on growth conditions^16,17^. The low CCE reflects the methane limiting conditions, where maintenance energy accounts for a higher proportion of methane oxidation. That a high respiratory demand decreases the growth efficiency has been shown for various bacteria in lakes^18^. According to the growth model, the MOB community indeed grew on average with only 26 ± 19 % (SD) of the maximum growth rate. In addition, the potentially higher metabolic costs of the elevated methane affinity required to thrive at low concentrations might also have reduced the CCE^13^. The best fit estimate for the maximum specific methane oxidation rate 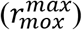 is 1.2 d^−1^ and lies well within the range of measured specific methane oxidation rates of 0.08 – 2.3 d^−1^ (ref.^12^). Combined with the carbon conversion efficiency the maximum specific methane oxidation rate translates to a growth rate of 0.16 d^−1^ or a doubling time of ~4 days. Doubling times reported for pure cultures are typically in the range of a few hours^13,14,19^, but these values represent growth under optimal conditions. Doubling times in environmental settings are likely to be slower and reported values for freshwater samples are in the range of 2 – 3 days^20–22^.

## Robustness of the microbial methane converter

The velocity of thermocline deepening and the piston velocity of gas exchange with the atmosphere, are important factors influencing the efficiency of the microbial methane converter (i.e. the amount of stored methane that is oxidized by MOB before it escapes to the atmosphere). During the autumn overturn in Lake Rotsee, wind and thermal convection played different roles in this respect. Our model calculations show that wind contributed 77 ± 13 % (SD) to the gas transfer coefficient and that wind was therefore the main driver for the transport of methane from the surface water to the atmosphere. In contrast, wind on average only contributed to 5% to the dissipation rate of turbulent kinetic energy at the mixed-layer depth (Figure 2a, inset) and the Monin-Obukhov length scale was smaller than the mixed layer depth (Supplementary Figure 5) for most of the time. Thermal convection was therefore the dominant driver for the deepening of the thermocline and thus for the supply of stored methane to the mixed layer. Model simulations of the autumn overturn in Lake Rotsee based on meteorological data from 1990 to 2016 showed that this convective regime dominated throughout the years. The annual variability of stored methane that is oxidized by MOB stayed in a fairly narrow range of 95% to 98%.

This high efficiency of the methane converter was a result of the convective mixing regime, with a gradual deepening of the thermocline that kept methane fluxes well below the methane oxidation capacity, defined here as the maximum methane oxidation rate at a given concentration of biomass (Figure 3a). The average of the daily maximum methane fluxes to the mixed layer reached only 27 % of the methane oxidation capacity even when deeper water layers with higher methane concentrations were involved. This robustness of the methane converter during the overturn is due to two effects. First, the increasing flux of stored methane to the mixed layer was diluted in an ever-larger volume of the mixed layer. The dilution of stored methane entering the epilimnion was three times larger at the end of the overturn compared to the initial situation. Second, the supply of methane led to an increase of MOB biomass, which in turn increased the methane oxidation capacity itself. In summary, both the progressing dilution of the methane supply as well as the growing methane oxidation capacity counteract the increasing flux of methane supplied to the mixed layer.

**Figure 3:**
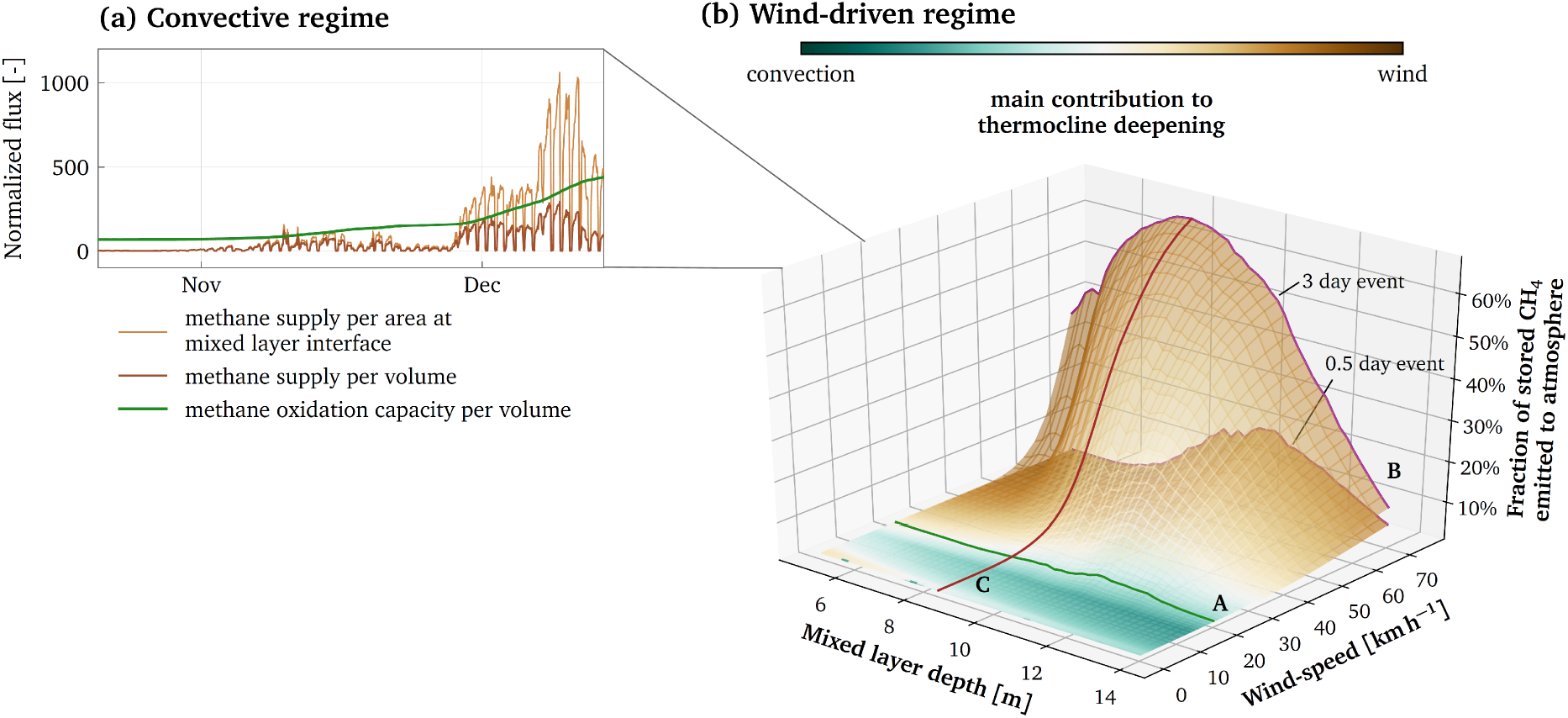
Robustness of the microbial methane converter. **(a)** Typical methane fluxes to the mixed layer for the convective regime. The methane fluxes were normalized to the minimum non-zero flux and are plotted per crosssection area at the mixed layer interface and per volume of the mixed layer. The flux of stored methane per volume can be compared to the methane oxidation capacity, the maximum rate of methane oxidation of the MOB biomass. **(b)** Scenario analysis for events of elevated wind-speeds. The two surfaces represent the fraction of stored methane that is emitted at the end of the overturn period if a 3 day or a 0.5 day wind-event is artificially generated during the overturn period. The two surfaces are colored according to whether wind or convection was the dominant driver for the thermocline deepening. Wind-events started at different mixed-layer depths and consisted of constant windspeeds for the whole duration of the event.

Because Lake Rotsee is wind-sheltered, we used a scenario analysis to explore the robustness of the microbial methane converter to events of high wind-speeds (Figure 3b). The scenario analysis revealed three important patterns. First, the fraction of stored methane that is emitted to the atmosphere by the end of the overturn period is negligible for wind-speeds of up to about 20 km/h (Figure 3b, line A). For these wind-speeds, convection still dominates the deepening of the thermocline. Second, for a 12 hour wind event, the fraction of emitted methane stayed below 20% even for wind-speeds of up to 70 km/h (Figure 3b, line B). To increase this fraction to 60%, 3 days of consecutive wind-speeds of 70 km/h were necessary. Third, the timing of the windevent matters. The fraction of emitted methane was highest when the wind-event occurred at intermediate mixed-layer depths (Figure 3b, line C) but was significantly lower if it occurred at the beginning or the end of the mixing period.

Accordingly, we suggest that global estimates of methane emissions from lakes should consider robust and effective methane oxidation even during overturn as the norm rather than as an exception. Wind has often been perceived as the strongest forcing for turbulent mixing^23^. In lakes however, convective processes are usually the dominant drivers for mixing and stratification because unlike wind-shear, they remain constant with decreasing surface area. Convective forces are therefore increasingly becoming recognized as important drivers for various biogeochemical processes in inland waters^23^.

We expect that future emissions of stored methane from temperate lakes remain low. Climate change will increasingly strengthen lake stratification and make partial mixing more common^24^. Even though more extensive anoxic conditions will allow more methane to accumulate, the slower mixing process resulting from strong stratification in turn facilitates its oxidation. In contrast, we expect that the relative importance of ebullition increases, especially when prolonged hypoxia increase the sediment methane concentration. Elevated methane emissions during autumn overturn are linked to exceptional meteorological events with persistently strong wind for several days or a combination of rapid cooling with strong winds. Estimates of the frequency of extreme wind and cooling events during the autumn overturn might therefore be particularly important to estimate current and future emissions of stored methane, since such events could have larger effects than the average warming trend^25^ itself.

## Methods

### Study site

Lake Rotsee is a small eutrophic subalpine lake in Switzerland that is 2.5 km long, 200 m wide and has a maximum depth of 16 m. The lake is oligomictic and exhibits a stable stratification from approximately May to September^26,27^. The fate of methane was determined during 7 field campaigns over the course of the autumn overturn from October to December 2016. Water samples were taken at the deepest point of Lake Rotsee at 47.0705° N and 8.3157° E.

### Methane concentrations

We measured methane concentrations in the water column using the headspace equilibration method. 20 mL water samples were filled into 40 mL serum vials. The serum vials had previously been prepared by addition of 4 g sodium hydroxide pellets, repeated flushing with nitrogen (N_2_) and evacuation to a final pressure of 600 mbar. We measured methane concentrations in the headspaces with a gas chromatograph (Agilent 6890N, USA) equipped with a Carboxen 1010 column (Supelco 10 m × 0.53 mm, USA) and flame ionisation detector. Samples that exceeded the calibration range were diluted with N_2_ and measured again. Dissolved methane concentrations were calculated according to Wiesenburg & Guinasso^28^. Control experiments showed that 4 g of solid NaOH produced 32.6 nmol methane on average, which was accounted for when calculating methane concentrations.

Methane concentrations in the sediment were determined in two duplicate sediment cores. Sediment cores were retrieved from the deepest part of Lake Rotsee with a UWITEC gravity corer (UWITEC, Mondsee, Austria). For methane sampling, predrilled PVC tubes of 6.3 cm 3 diameter and 60 cm length with holes of 1.2 cm diameter spaced at 1 cm were used. 2 cm^3^ aliquots of sediment were taken for methane analysis with syringes with cut-off tips by pushing the syringe through the taped holes. The sediment aliquots were directly injected into 113.7 ml flasks pre-filled with 7 M NaOH, capped and stored in the dark at 4°C. Methane was determined by headspace analysis using an Agilent gas chromatograph (Agilent Technologies AG, Basel, Switzerland) equipped with a Supelco Carboxene®-1010 column (Sigma-Aldrich, Steinheim, Germany).

### Biomass of methane oxidizing bacteria

We determined MOB cell numbers using CARD FISH and epifluorescence microscopy. Water samples of 5 mL were fixed with 300 μl sterile filtered (0.2 μm) formaldehyde (2.22% [v/v] final concentration) for 3 – 6 h on ice. Samples were then filtered onto 0.2 μm nuclepore track-etched polycarbonate membrane filters (Whatman, UK), dried, and stored at –20°C until further analyses. Cells were permeabilized with lysozyme (10 mg mL^−1^) at 37 °C for 70 min, followed by inactivation of endogenous peroxidases with 0.1 M hydrogen chloride for 10 min at room temperature. Filters were hybridised with hybridisation buffer containing HRP-labelled probes at 46 °C for 2.5 h. The buffer contained either a 1:1:1 mix of Mg84, Mg705, and Mg669 probes targeting methanotrophic gammaproteobacteria or a Ma450 probe targeting methanotrophic alphaproteobacteria^29^. Fluorescent signals were amplified with the green-fluorescent Oregon Green 488 tyramide (OG) fluorochrome (1 μl mL^−1^) at 37 °C for 30 min. Hybridised cells were counterstained with DAPI (20 μl of 1 μg mL^−1^ per filter) for 5 minutes. We used a 4:1 mix of Citifluor and Vectashield to mount filter pieces for fluorescence microscopy. We quantified MOB cell numbers using an inverted light microscope (Leica DMI6000 B, Germany) at a 1000-fold magnification. For each sample, 22 image pairs (DAPI and OG) of randomly selected fields of view (FOVs) were taken for each filter piece. Cells were detected and counted using Daime 2.0 for computer digital image processing (CDIP)^30^. We converted the cell numbers to biomass using an average cellular carbon content of 0.42 pg cell^-1^ based on literature values^21,31,32^.

### Methane emissions

Methane emissions during the overturn period in 2016 were quantified with eddy-covariance flux measurements using an FGGA-100 fast methane and CO_2_ gas analyzer (Los Gatos Reseach Inc., Mountain View, CA, USA) combined with a Solent R2A ultrasonic anemometer (Gill Instruments, Solent, UK) as described in Sollberger et al.^33^ and Eugster & Plüss^34^. The measurement setup was deployed at the same location as the one described in Schubert et al.^8^: The sonic anemometer was mounted on a moored buoy 70 m from the lake shore, and air was pulled 1. 47 m (center of ultrasonic anemometer head) above the lake surface and guided to the gas analyser using a Synflex 1300 tube at a flow rate of 26.3 L min^-1^. The footprint area of the eddy covariance measurements is shown in the Supplementary Figure 3.

### Meteorological conditions and water temperature

We continuously measured the evolution of the water temperature with a temporal resolution of 5 s and a spatial resolution of 1 m (RBRconcerto T24 thermistor string, RBR). To be able to cover a long period, air temperature, wind speed, vapour pressure, shortwave incoming radiation and cloud cover were obtained from MeteoSchweiz (SwissMetNet station LUZ). We adjusted wind-speeds to the local conditions using sporadically available wind speed measurements directly at Lake Rotsee obtained with an automated weather station by AANDERAA (Figure S1).

### Biogeochemical model

The mixed layer was approximated as a single, homogeneously mixed box, which gets larger as the thermocline deepens. The system of ordinary differential equations (ODE) describing the temporal evolution of the mixed-layer depth (*h_mix_*) and volume (*V_mix_*), temperature (*T_mix_*), methane concentration (*C_mix_*) and MOB biomass (*B_mix_*) is:

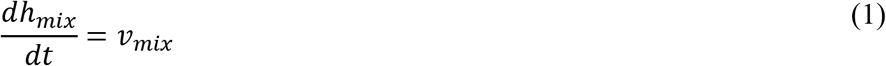

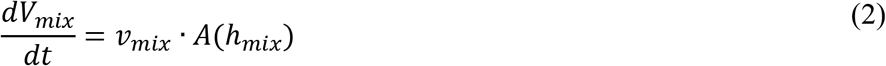

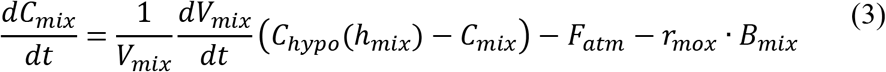

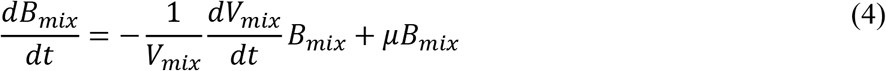

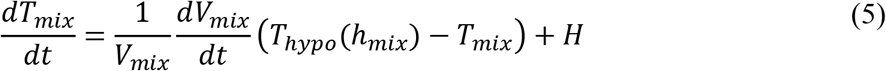

where *v_mix_* is the velocity of the thermocline deepening, *A*(*h_mix_*) is the cross-section area of the lake at the mixed-layer depth *h_mix_, F_atm_* is the flux of methane to the atmosphere, *r_mox_* is the specific rate of microbial methane oxidation, *μ* is the growth rate of MOB, *H* is the heat flux into (positive) the lake, and *C_hypo_*(*h_mix_*) and *T_hypo_*(*h_mix_*) are the methane concentration and temperature in the hypolimnion at the mixed-layer depth.

The initial temperature profile at the onset of the overturn period was obtained from field data and was assumed to remain constant in the hypolimnion. In contrast, the measured methane profiles do not remain constant due to the ongoing accumulation of methane at the bottom of the lake. To keep the model simple, instead of modelling the accumulation we compiled a profile of maximum concentrations (Supplementary Figure 2) that was used as the initial methane profile. In the above system of ODEs, *C_hypo_* and *T_hypo_* are only depth dependent and correspond to the temperature and methane concentration given by the initial profile at the mixed-layer depth.

We calculated methane emissions with the boundary layer model of Liss and Slater^35^:

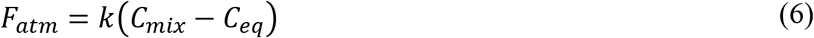

where *C_eq_* is the water saturation concentration calculated according to ref. ^36^ and *k* is the transfer velocity. To explicitly take into account buoyancy and wind driven components of turbulence at the air-water interface, we calculated the transfer velocity using the surface-renewal model^37–39^:

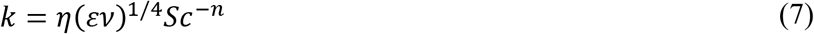

where *ε* is the rate of dissipation of turbulent kinetic energy near the air-water interface, *v* is the kinematic viscosity and *η* is an empirically derived, depth dependent scaling coefficient. For the purpose of this conceptual model, *η* was set to 1/(2π) based on theoretical considerations^40^. We determined the Schmidt number *Sc* for methane with an exponent *n* = −1/2 (ref. ^36^).

The dissipation rate *ε* was explicitly partitioned into the wind shear and the buoyancy components:

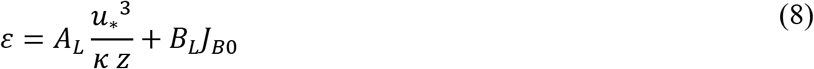

where *k* is the von Kármán constant, *u*_*_. is the water-side friction velocity and *A_L_* as well as *B_L_* are empirically derived coefficients set to 1 as in ref.^39^. The wind-shear component of the dissipation was calculated at the depth of the viscous boundary layer *z* = *δv* (ref. ^40^):

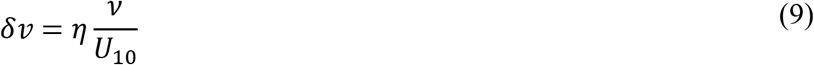

We calculated the friction velocity *u*_*_. as a function of the 10-m wind speed *U*_10_ as described in ref.^41^:

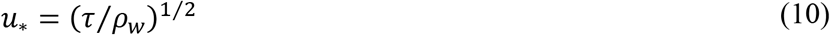

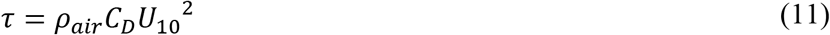

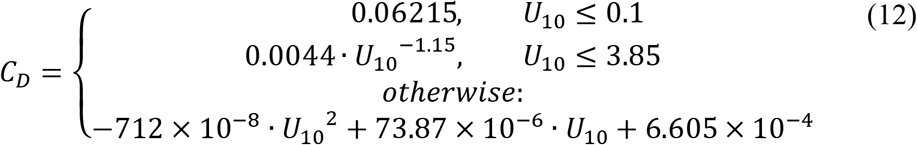

The buoyancy flux is defined as: *J_B0_* = *gαH*/(*p_w_C_p_*) with the gravitational constant *g*, the thermal expansion coefficient *a,* the heat flux *H*, the density of water *p_w_* and the isobaric heat capacity *C_p_*. Finally, the parameterization of the heat flux *(H)* includes the shortwave absorption, the longwave absorption, the longwave emission, the heat flux by evaporation and the heat convection^42^.

The deepening of the mixed layer was parameterized as a function of convective and wind-driven forcing^43,44^:

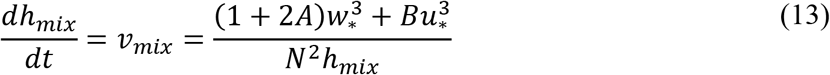

where A and B are respectively entrainment coefficients for the convective, *w*_*_, = (*J_B0_ h_max_*)^1/3^, and sheared *u*, velocities and *N*^2^ is the Brunt-Väisälä frequency at the mixed-layer interface^45^. We set *A* = 0.2 (ref. ^46^) and *B* = 2.5 (ref. ^47^).

In the mixed layer, we assume that methane is the limiting nutrient and we formulate the growth rate of MOB with a Monod-type methane oxidation kinetics where the growth rate (μ) is approximated based on the maximum specific methane oxidation rate *(r_mox_*) and a carbon conversion efficiency (*y*):

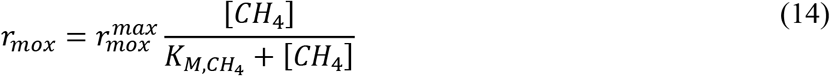

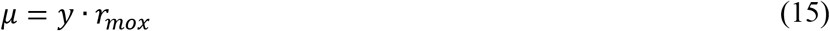

We obtained the kinetic parameters 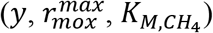 based on the measured biomass evolution. First, we performed a global parameter estimation using the CRS algorithm^48^. Based on the result, a local parameter estimation was performed using the COBYLA algorithm^49^.

The system of ODEs was implemented with Julia 1.0.1^50^ and solved numerically with adaptive time stepping and a maximum time step of 1 hour using an order 2/3 L-Stable Rosenbrock-W method – which is good for very stiff equations with oscillations at low tolerances – provided in the *DifferentialEquations.jl* package^51^. Parameter estimation was performed using the Julia package *NLopt.jl*^52^.

### Fate of Methane

We calculated the methane budget neglecting potential horizontal variability. Amounts of methane and biomass in the water column are given as volume integrated mass of carbon. We obtained profiles of methane and biomass storage in units of Mg m^−1^ by multiplying the measured concentrations with the cross-section area of the lake at the respective depth of measurement. Methane emissions were extrapolated to the whole lake surface area and are given as cumulative amounts.

The statistical interpretation of number ranges is given in brackets. We used the standard deviation (SD) when the variability of the sample was of interest. The standard error of the mean (SEM) or the 95% confidence interval (CI) was used in cases where the uncertainty of the mean was of interest. The confidence intervals were corrected for the sample size using the t statistic.

### Robustness of the microbial methane converter

The contribution of wind to the gas transfer coefficient was expressed as:

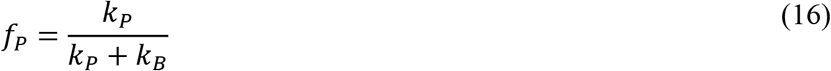

where *k_P_* is the transfer coefficient based on wind and *k_B_* is the transfer coefficient based on convection.

Because Lake Rotsee is wind-shelterd, we explored the effect of various wind regimes on methane emissions by a model scenario analysis. We set wind-speeds to a constant level for 12 hours or 3 days. Each scenario consisted of a single wind-event that started at a specific mixed-layer depths. To avoid artefacts from other meteorological conditions during the wind-event, we neglected the heat balance during this period and calculated the forced convection based on an empirical regression between the friction velocity and the buoyancy flux at the surface (Supplementary Figure 4).

## Code availability

The source code of the generic model is available as a Julia package on GitHub: https://github.com/zimmermm/ConvectionBoxmodel.jl. The actual implementation for Lake Rotsee is attached to the data package on the ETH Research Collection (10.3929/ethz-b-000350091).

## Data availability

All measurement data used for this publication are available on the ETH Research Collection (10.3929/ethz-b-000350091).

Meteorological data was obtained from MeteoSwiss (www.meteoswiss.admin.ch)

## Acknowledgments

This research was supported by SNF (grant CR23I3_156759) and ETH Zurich (scientific equipment grants 0-43350-07, 0-23138-09 and 0-43683-11). We thank Karin Beck, Michael Plüss, Michael Schurter and Christian Dinkel for the technical support during the field campaigns and Jason Dey for CARD-FISH and image analysis.

## Notes

http://doi.org/10.3929/ethz-b-000350091

