## Supporting information for "Lake overturn as a key driver for methane oxidation"

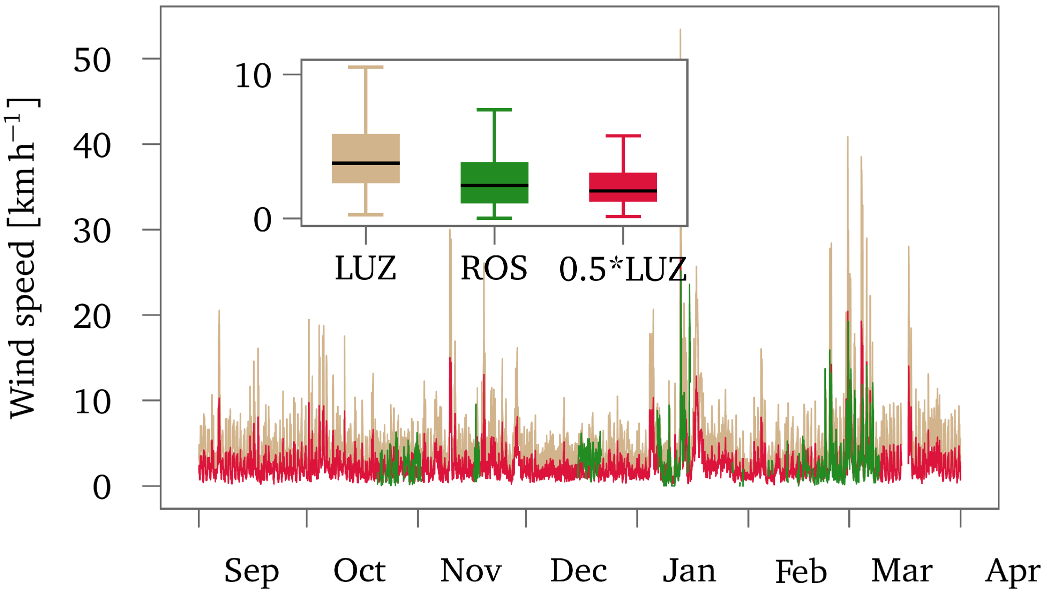


**Supplementary Figure 1**. Inference of wind speeds at Lake Rotsee from a nearby automated weather station. Wind speeds measured by an automated weather station by MeteoSwiss on a nearby hill (LUZ) are plotted in gray. Sporadically available wind speed measurement directly at Lake Rotsee (ROS) are plotted in black. We scaled the wind-speeds of the weather station by MeteoSwiss by 0.5 to get continuous wind speeds that are comparable to wind speeds measured directly at Lake Rotsee (plotted in red).

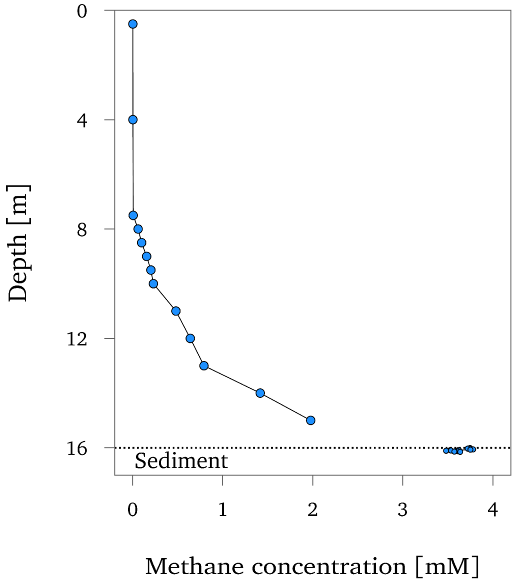


**Supplementary Figure 2**. Profile of maximum methane concentrations in Lake Rotsee. Each dot represents the maximum concentration measured during the overturn period in the respective depth.

4

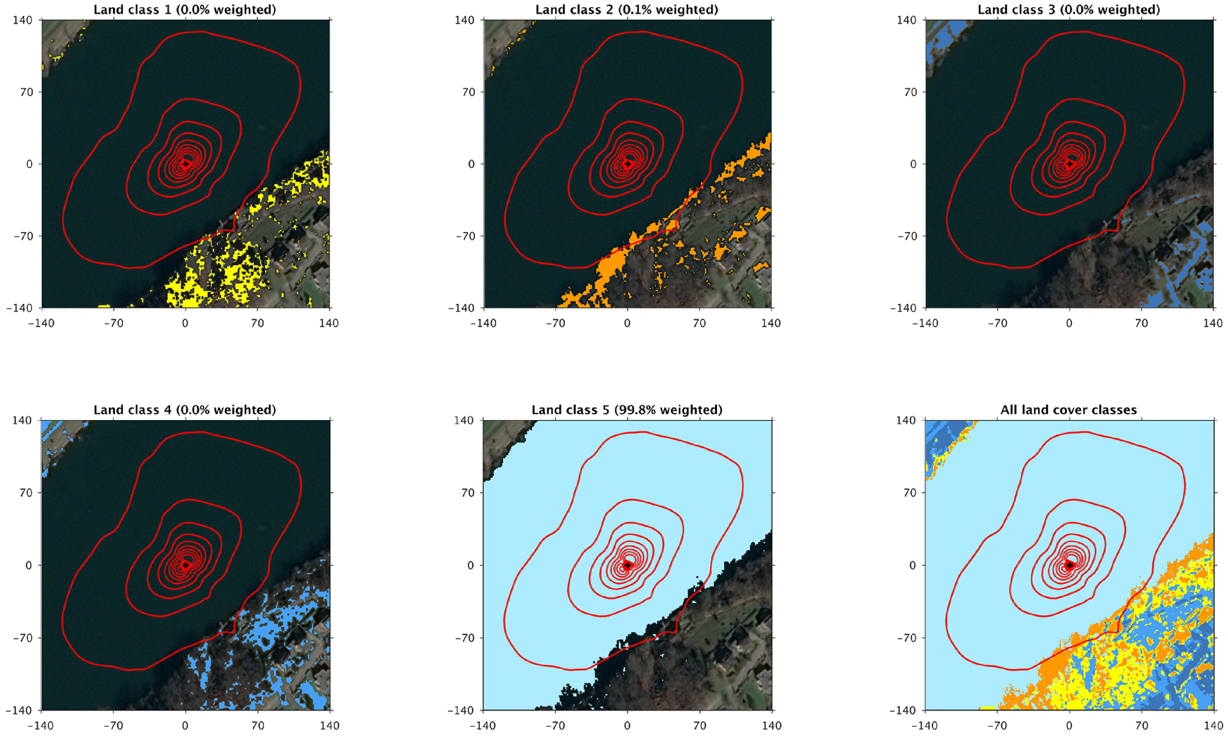


**Supplementary Figure 3**. Footprint of the eddy flux measurement. Google map of the flux tower area with footprint contour lines from 10 to 90%, in 10% steps for five unsupervised land cover classifications. Footprint calculation and land cover classification were done using the FFPonline tool (http://footprint.kljun.net/ffp2d.html).

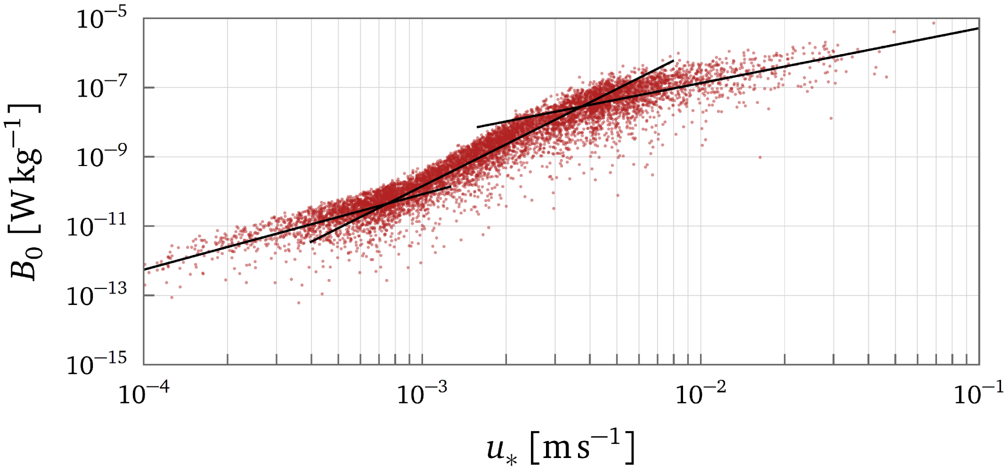


**Supplementary Figure 4**. Correlation between friction velocity and surface buoyancy flux during lake overturn in autumn 2016 in Lake Rotsee.

5

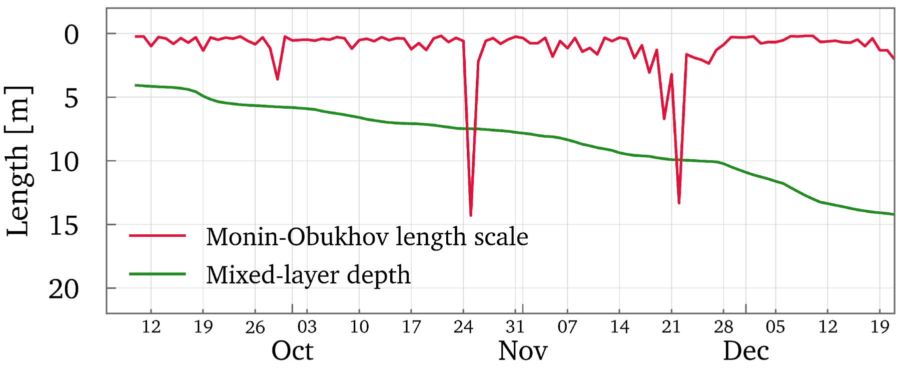


**Supplementary Figure 5**. Daily average Monin-Obukhov length scale and mixed-layer depth. Convection dominates the thermocline deepening if the Monin-Obukhov length is smaller than the mixed-layer depth. Otherwise, wind dominates the mixing process.
